# Predicting Cerebral Pericyte Contractility Across Experimental and Physiological Conditions: an in-silico framework

**DOI:** 10.64898/2026.09.01.748462

**Authors:** Alberto Coccarelli, Ammar Al-Areqi, Osama F. Harraz

## Abstract

Pericytes (PCs) have recently emerged as critical regulators of cerebral blood flow (CBF) and represent a promising therapeutic target for various cerebrovascular pathologies. Given the complex array of biochemical and mechanical stimuli these cells integrate, a multiscale modeling framework is essential to quantify the impact of selective interventions on pericyte contractile machinery and blood flow restoration. Here, we introduce a computational framework to evaluate capillary pericyte responses across diverse experimental interventions and conditions (ex vivo and in vivo). To capture pharmacological modulation of the contractile apparatus, we developed a homogeneous intracellular model that incorporates key properties of robust control systems. In this framework, vascular tone generation depends strictly on intracellular calcium concentration (Ca^2+^), which emerges from a complex electrochemical equilibrium established by transmembrane ion (Na^+^, K^+^, Cl*^−^*) gradients, luminal mechanical forces, and external ligand concentrations. The resulting fraction of phosphorylated cross-bridges generates contractility, which is integrated into the strain energy function governing the constitutive behavior of the vascular wall. The model was successfully validated across four distinct experimental and pharmacological interventions (including pinacidil, high external K^+^, U46619, and nimodipine), demonstrating close agreement with observed ex vivo and in vivo vascular responses. By establishing a quantitative bridge between pericyte electrophysiology and microvascular mechanics, this framework provides a valuable foundation for evaluating targeted therapeutic strategies to alleviate tissue ischemia in stroke and vascular dementia.

## 1 Introduction

Compromised cerebral blood flow (CBF) is implicated in various pathological conditions, including vascular dementia and stroke. In principle, brain perfusion can be improved using vasodilators that selectively relax the musculature of these tiny vessels, enabling more blood flow and oxygen/nutrient delivery to surrounding tissue. However, these therapeutic approaches are hindered because the microvasculature comprises functionally distinct regions that are haemodynamically and electrically coupled, making it difficult to isolate their individual contributions and identify optimal vascular compartments to target. Over the past decade, pericytes (PCs) have emerged as key regulators of CBF. Positioned at the transition between arteriolar and deep capillary regions, PCs modulate a major fraction of cerebrovascular resistance [21, 19, 17, 24, 14]. These cells exhibit functional properties distinct from smooth muscle cells (SMCs), which also vary along the arteriolar-capillary transition (ACT) zone [20]. Their strategic position requires them to regulate CBF within a complex microenvironment, integrating diverse biochemical signals from neighboring neurons, glia, and endothelial cells alongside mechanical forces and circulating hormones. Experimental characterization of pericyte function in vivo is challenged by their dense microenvironmental integration, limited accessibility, and multi-scale spatiotemporal dynamics. Furthermore, despite the central role of calcium signaling in tuning cerebral vascular tone [4], Ca^2+^ signatures vary considerably across the microvascular bed [3, 16, 31, 41]. Consequently, available experimental literature remains fragmented and difficult to compare due to variations in protocol, measurement modality, and experimental focus. Recent mathematical models have begun characterizing specific aspects of rodent cerebral microvascular function, such as retrograde hyperpolarization signal propagation through the endothelium [40] and differences in myogenic tone between arteriolar SMCs and PCs [30]. However, sparse experimental evidence has hindered the development of identifiable, highly generalizable multi-scale models. To date, no unified computational framework seamlessly integrates pericyte data across diverse experimental setups while validating predictions against multiple pharmacological interventions. Establishing such a platform is essential for translating new or repurposed vasoactive agents into clinical applications. In this study, we present a computational modeling framework capable of accurately capturing ex vivo experimental data and generating predictive insights across distinct pharmacological scenarios. This framework provides a foundation for quantifying modulation thresholds in interventions targeting the cerebral microvascular contractile machinery.

## 2 Modelling framework

Our model specifically represents the contractile mural of the 1st-3rd order of capillaries in the ACT zone, the so called ensheating pericytes. Upon some modifications, the model can be used to mimic the contractility of deeper capillary pericytes which substantially differ from the upstream ones.

### 2.1 Main assumptions

The main assumptions underlying cell’s contractility and deformation are:

- the cell possesses features typical of a robust control systems, for which physiological stimuli shifts the mechano-electrochemical equilibrium from one calcium steady state to another, which can be encoded by wholecell mean (time-averaged) intracellular Ca^2+^, while features such as peak amplitudes and oscillation frequencies are disregarded;
- channels, pumps, exchangers, and receptors, as well as chemo-mechanical stimuli, are homogeneously distributed along the cell membrane. Unless specified otherwise, ion channel conductances and cross-bridge (XB) dynamics depend solely on the averaged cytosolic effector concentrations;
- the model captures primarily only short-to-medium (*≈* minutes) vascular adaptation responses, whereas long-term structural remodeling processes, such as cytoskeletal rearrangement and immuno-mechanical pathways, are beyond the scope of this work;
- the cell structure is represented as a deformable ring wrapping around the whole circumference of the capillary, with no functional distinction between soma and processes. In this study only the mechanics along the radial direction is considered but the cell axial length could scaled to resemble the effective PC vessel coverage and its heterogeneous spatial variability;
- the surrounding neurons, endothelial and glial cells are not explicitly represented but their contribution to tone regulation can be accounted for by modulating the membrane potential and ion channels’ activity.

### 2.2 Cell electrophysiology

External stimuli such as luminal pressure/wall stress (*σ*) and vaso-active agents modulate the PC active tone by changing membrane protein activity and triggering intracellular signalling pathways which ultimately redistribute the fraction of phosphorylated XBs promoting the acto-myosin filament sliding. Components of the proposed model architecture (Fig. 1) are derived from previous computational studies on SMCs and PCs [52, 28, 8, 30] and is informed by recent experimental evidence [48, 33, 31, 32, 47, 12]. Here the considered electrophysiological variables are the membrane potential (*V*_m_), the spatially-averaged intracellular concentration of Ca^2+^, Na^+^ and Cl*^−^* (Ca_i_, Na_i_ and Cl_i_), and the store Ca^2+^ concentration (Ca_s_).

**Figure 1:**
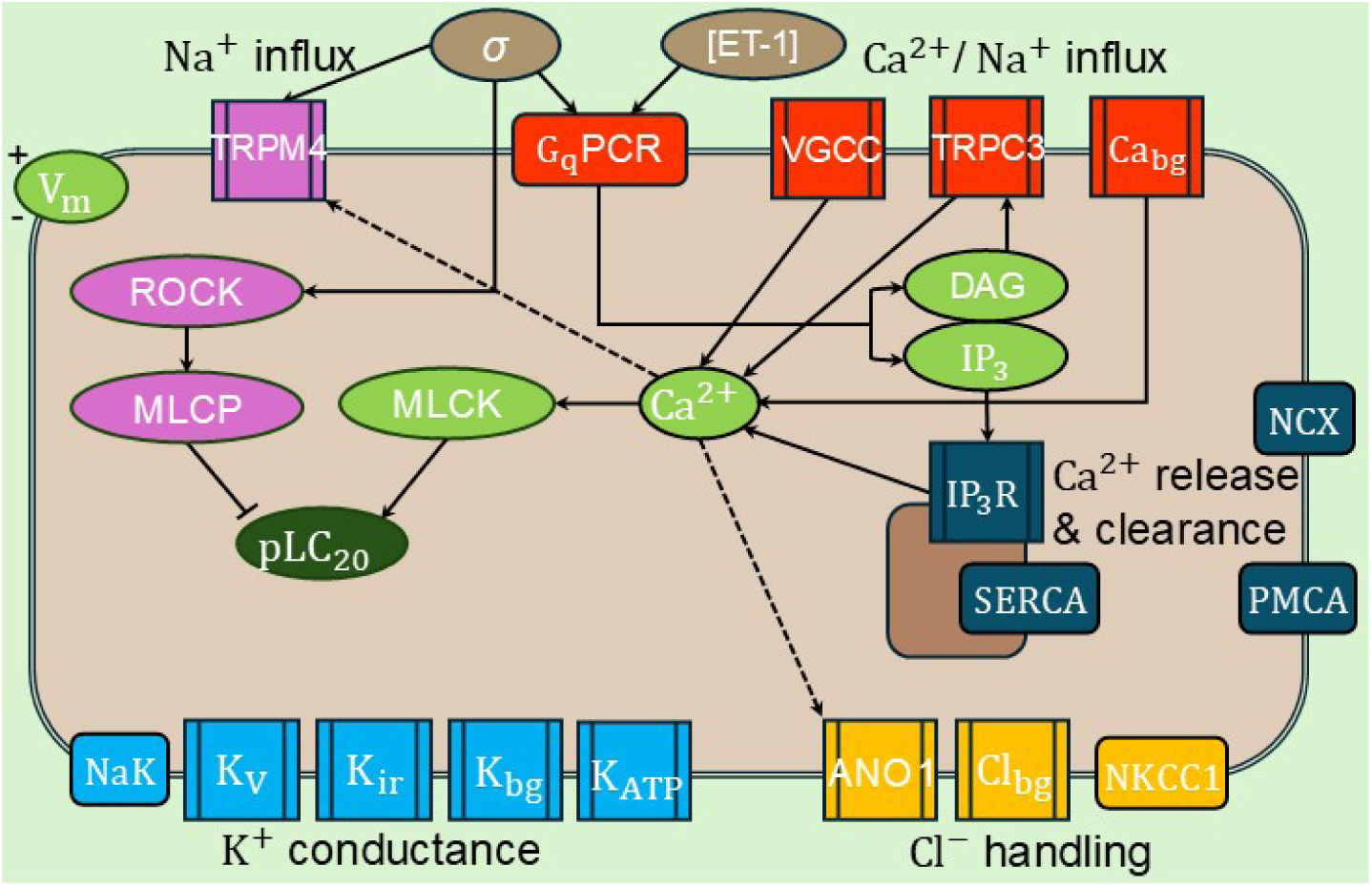
Schematic representation of the PC contractility model, with focus on the associated Ca^2+^ signalling. Black arrows indicate activation pathways (dashed pattern associated with Ca^2+^activated pathways). pLC_20_: phosphorylated myosin light chain 20.

#### 2.2.1 GPCRs activation and Cation influx

There is compelling evidence that G-Protein-Coupled Receptors (GPCRs) are primary sensors of fluid forces mediating myogenic tone across different types of microvessels [43, 39, 29, 7]. In the context of cerebral pericytes, the review by Hariharan et al. [22] provided, by leveraging data from previous brain single-cell RNAseq screens, an exhaustive summary about the nexus between GPCRs and ion channels. In this study, the G_q_-coupled receptors (G_q_PCRs) are modelled as key mediators of mechanosensation and ligand-induced contraction, transducing the effect of primary vasoactive agents such as endothelin-1 (ET-1), angiotnesin II (Ang II) and thromboxane A_2_ (TXA_2_). According to the current paradigm, stress on the capillary wall leads to an increase in the local production of inositol 1,4,5-trisphosphate (IP_3_) and diacylglycerol (DAG), activating the nearby nonselective cation channels, which depolarize the cell [25, 6]. A recent study [12] identified transient receptor potential canonical 3 (TRPC3) as a primary driver for pressure-induced tone in cerebral pericytes. It has also been reported that wall stretch enables angiotensin II type 1 receptor (AT_1_R)-TRPC3 coupling by an alternative biased signalling [37], enhancing G_q_PCR activation downstream effects (with respect to ligand-induced activation alone), as reported in a recent computational study [42]. Previous studies [13, 35, 2] reported an elegant and effective methodology to model GPCR activation and subsequent increase in intracellular IP_3_. Due to the limited availability of pericyte-specific data, the dynamics of GPCRs activation and subsequent phospholipase C (PLC) signalling are not explicitly modelled; instead, their activation is directly represented as an intracellular increase in IP_3_ and DAG. The resulting cytosolic level of IP_3_, which is sensed by the IP_3_Rs, is assumed to be given as the sum of a ligand (such as [ET-1]) and stress dependent (*σ*) G_q_PCR contributors, which are modelled via Hill activation functions:

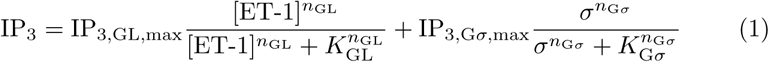

where IP_3,GL,max_ and IP_3,G*σ,*max_ are the corresponding maximum IP_3_ concentrations, while *n*_GL_*, K*_GL_*, n*_G*σ*_*, K*_G*σ*_ are the Hill function coefficients. The first term on the RHS of Eq. (1) represents the widespread sustained elevation in IP_3_ across the cytosol due to classical ligand-activation of the G_q_PCRs, while the latter term reflects an additional IP_3_ booster associated with the pressure-induced biased signalling.

The increase in TRPC3 conductivity leads to considerably net influx of Na^+^ and Ca^2+^, depolarizing the cell and modulating the activity of multiple membrane proteins. Klug et al. [31] showed that upon pressure elevation, voltage gated calcium channels (VGCCs) act as the dominant Ca^2+^ influx the first 3-4 capillary generations. This is accompanied by TRPC, calcium background and Orai currents, with the last likely being a contributor in the deep capillary regions [41]. The TRPC3, VGCC and Ca^2+^-background currents, elevating intracellular calcium upon pressure elevation and/or membrane depolarization are described via GHK formalism (see Supplementary Information, Section S3). Despite it has been reported that in cerebral SMCs TRPC3 can be directly activated by IP_3_Rs through physical coupling [1], here we assume for simplicity that TRPC3 are activated solely by the wall stress-generated DAG (through G_q_PCR), whose activation function coincides with 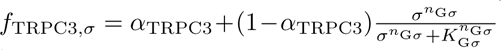 where *α*_TRPC3_ is the basal stress-independent fraction. VGCCs subtypes are not modelled distinctively but its voltage dependency is scaled to account for all the net contributors. The background calcium channels include an Orai/store-depletion component as done in [28] and present a rectification term to limit the influx current at very low *V*_m_. mRNA data [49] suggest that, due to its high expression, transient receptor potential melastatin 4 (TRPM4) may also contribute to myogenic tone in the ACT zone, which is in line with the study by Gonzales et al. [18] that characterized stretch dependent activation of TRPM4 via AT_1_R in cerebral vessels. Data from [12] also suggests that there might be at least another depolarizing current in PCs when pressure increases, but to date there is not clear evidence on the origin. To reflect this, we introduce an extra pressure-dependent current (representing potential TRPM4 contribution) which is only Na^+^-conductive, and activated by intracellular Ca^2+^. The classical Ca^2+^ extrusion and sequestration mechanisms via plasma membrane Ca^2+^-ATPase (PMCA), Na^+^*/*Ca^2+^ exchanger (NCX) and sarcoendoplasmic reticulum Ca^2+^-ATPase (SERCA) are described using reference formulations [52, 28].

#### 2.2.2 Calcium store release and membrane voltage regulation

Increase in cytosolic IP_3_ trigger calcium release from stores (only IP_3_Rs are considered), depleting calcium reservoir and enhancing calcium influx through Orai channels. Given the limited data on PC IP_3_ flux dependency on calcium and IP_3_, we used a formulation in line with reference work [2, 28]. Notably, the shape of inositol 1,4,5-trisphosphate receptor (IP_3_R) flux (*J*_IP3R_) vs Ca_i_ plays a negligible role at steady state as it is balanced out by leak and SERCA sequestration. Calcium release from sarcoplasmic reticulum (SR) has twofold effects: increase temporarily the intracellular calcium and promote opening of the calcium-activated Chloride channels ANO1 (TMEM16A), which further depolarizes the cell membrane. Since these channels are in close contact with IP_3_Rs forming Ca^2+^ micro-domains, their calcium-depended activation is modelled as a Hill function of whole cell and localized IP_3_Rs flux:

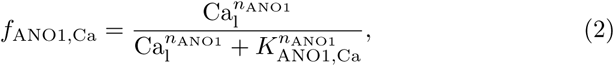

where Ca_l_ = Ca_i_ + *β*_ANO1_*J*_IP3R_, with *β*_ANO1_ being a scaling factor converting flux into local concentration augmentation. The ANO1 Cl*^−^* efflux is balanced out by the Na-K-Cl cotransporter (NKCC1) and Cl*^−^* background current. The established IP_3_ *− f*_ANO1,Ca_ dependency enables strong membrane depolarization upon ligand-G_q_PCR activation, and non negligible contribution to *V*_m_ when pressure increases. The NKCC1 is endowed with an inactivation term that limits the transporter conductance for increasing intracellular Cl*^−^*. It is very likely that other proteins regulate Cl*^−^* homeostasis but the limited experimental characterization of PCs prevented us to include other distinct components. The steady-state membrane potential is set as a result of the interaction between the above mentioned channels and potassium channels, which have been characterized in cerebral PCs by Sancho et al. [48, 47]. Here voltage-gated, inward rectifier, background and ATP channels are considered, with the latter only active upon selective stimulation (such as Pinacil application) since ATP dynamics is not included in the current modelling framework. K^+^ efflux (and Na^+^ influx) is primarily counterbalance by the NaK exchanger, whose dependency on Na^+^ and external potassium (K_o_) is dealt via double Hill function [28]. Since the intracellular K^+^ is expected to remain extremely high (140 mM) upon the vast majority of physiological scenarios, we assume it to remain constant over time and a Ohmic approach is used to describe the K_v_, K_ir_, K_Bg_ and K_ATP_ currents. The membrane voltage and ion mass balances, alongside the current definitions are reported in the Supplementary Information Section S3.

### 2.3 Capillary deformation

Experimental evidence [24] suggests that, alongside GPCR activation, pressure may activate calcium sensitization pathways, such as Rho-associated protein kinase (ROCK), in line with findings from previous studies in cerebral arteries [34]. Herein is assumed that the PC active tone is solely driven by the myosin-actin XBs dynamics, which is made dependent on the intracellular calcium dynamics and calcium sensitization processes (ROCK activation), as done in our previous studies on cerebral arteries myogenic regulation [8, 9]. The resulting intracellular calcium concentration is dictated by the combination mechanical forces (wall stress/pressure) and external ligand concentration (ET-1 and channel activators/inhibitors), and determines the level of myosin light chain kinase (MLCK) activation:

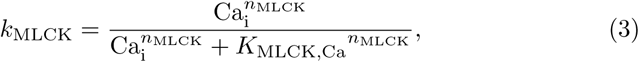

where *k*_MLCK_ is the (normalized) MLCK-induced phosphorylation rate, while *K*_MLCK,Ca_ and *n*_MLCK_ are the Hill function parameters linking the MLCK activity with the intracellular calcium level. In cerebral SMCs, purinergic receptors are tightly associated with Rho-kinase activity, which can also activate TRPM4 [36], suppress K_v_ currents [38], promoting cell depolarization. It is also possible that pressure modulates the activity of K_v_ and other ion channels by altering phosphatidylinositol 4,5-bisphosphate (PIP_2_) regulation [44, 15]. Nevertheless, given the limited experimental evidence on this, we simply assume that wall stress contributes to tone via a secondary pathway, which involves ROCK activity increase with consequent reduction of myosing light chain phosphatase (MLCP) capacity to de-phosphorylate XBs [8].

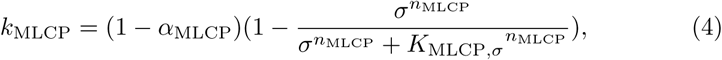

where *k*_MLCP_ is the (normalized) MLCP-induced dephosphorylation rate, *α*_MLCP_ is the basal stress-independent component, while *K*_MLCP,*σ*_ and *n*_MLCP_ are the Hill function parameters representing the dependence of ROCK activity on wall stress. The activated (phosphorylated) XB fraction (*n*_AMp_) is evaluated via the mass conservation balance:

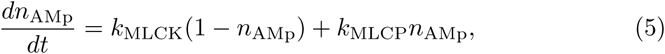

As in our previous works [8, 9]:

- the tone generated by the phosphorylated XBs is encoded in the active component of the strain energy function, which reflects the constitutive behaviour of the vascular wall;
- the passive component of the vascular response is dealt with an incompressible hyper-elastic formulation;
- vascular tone and deformation are evaluated by ensuring that the momentum conservation equation along the ring radius is satisfied.

With respect to our previous work [8] the forces governing the sliding filament kinetics (*F*_XB,a_ extracellular matrix reaction, *F*_XB,c_ power stroke) are simplified as:

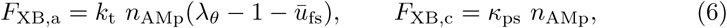

where *k*_t_ is the stiffness associated with XBs deformation, *n*_AMp_ is the phosphorylated XBs fraction, *λ_θ_* is the circumferential stretch, *u̅*_fs_ is the (normalized) relative filament sliding (variable) and *κ*_ps_ is the power-stroke magnitude per phosphorylated XB. Given the limited vessel thickness, we assume that the intracellular signalling depends only on the luminal wall stress (pressure) and that does not vary along the thickness of the vascular ring.

### 2.4 Parameters identification strategy

The positive (regenerative) feedback loop driving Ca^2+^ elevation provides the basis for bistability, potentially conferring a switch on-off behaviour to the model. Provided this, the choice of parameters dictates whether the cell behaves as monostable or bistable system. In this study, we define a strategy such that the developed PC contractility model possesses, to some degree, the features of a robust control system, such that it produces a gradual response to various mechano-chemical stimuli, over a realistic range of conditions. To achieve this the intracellular signalling model is parametrized through two steps. First, by using whole-cell patch clamp data, we define priors for different ion channels parameters. We use the datasets from [53, 33, 47, 12, 30] for informing the currents of K_ir_, K_v_, TRPC3, VGCC and ANO1 channels. Then we identify the key model parameters (30) by solving a ‘cellular signalling’ optimization problem for which the objective function aims to satisfy a set of constraints on PC steady state solutions upon different types of stimulation (such as pressure increase and drug modulating channel activity) and conditions. The first subset of constrains is defined based on experimental rodent data from two ex-vivo protocols, using a pressurized cannulated microvascular (CaPA) network with no flow and superfused acute brain slices (Table 1). For some target variables (such as *V*_m_ at rest, *≈* at 10 mmHg with no drugs) we opted to define a relatively broad target range of values to reflect the variability observed across experimental studies [48, 31, 12], which might also be significantly affected in vivo by the activity of the surrounding cells [21, 23]. Ca_i_ variation (with respect to baseline) is converted to relative fluorescence variation via 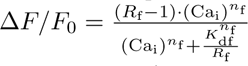 where *R*_f_ is the dynamic range of the indicator (*F*_max_*/F*_0_), representing the ratio of maximum to minimum fluorescence, which is kept constant throughout all simulations. The parameters *K*_df_ and *n*_f_ are indicator-dependent, evaluated here for the genetically encoded calcium indicators GCaMP5G and GCaMP6f, and are taken from [5].

**Table 1:**
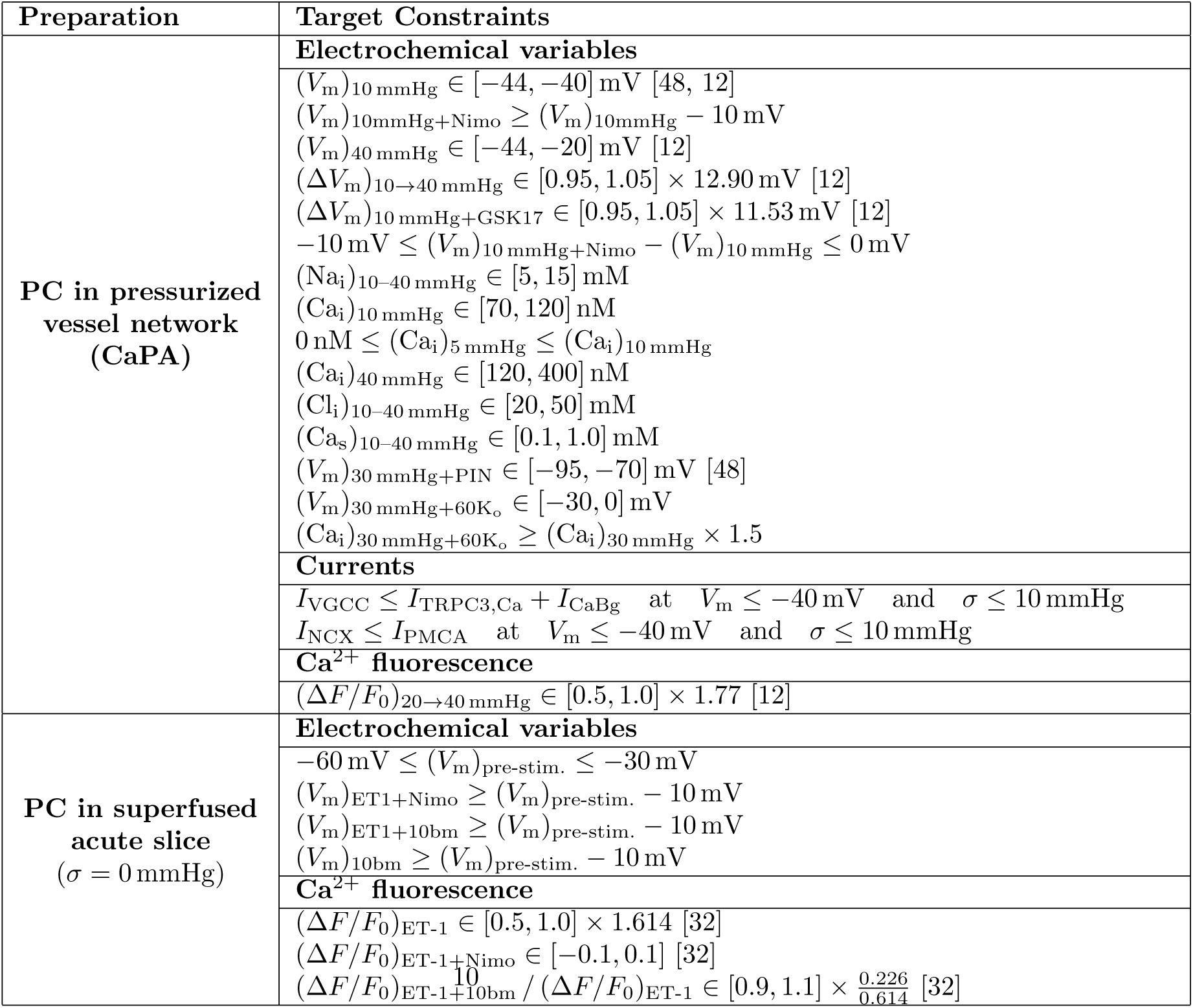
Physiological constraints used in the model parameter identification. For CaPA preparation: Na_o_=152.25 mM, K_o_=3 mM, Ca_o_=2 mM, Cl_o_=134 mM. For acute slices preparation: Na_o_=151 mM, K_o_=2.5 mM, Ca_o_=2 mM, Cl_o_=132.5 mM. GSK1702934A (GSK17, 100 nM) is expected to fullyopen TRPC3 channels, for which *f*_TRPC3*,σ*_ = 0.9 is assumed. Nimodipine (Nimo, 3 *µ*M) is assumed to reduce VGCC’s permeability-surface area product (*PA*_m_)_VGCC_ by 90%. Similarly, the ANO1-blocker 10bm (200 nM) is assumed to reduce ANO1’s permeability-surface area product (*PA*_m_)_ANO1_ by 90%. Application of Pinacidil (PIN, 10 *µ*M) activates K_ATP_ channels, causing a strong cell hyperpolarization. Based on intact whole-cell current recordings in [48], we estimated a (maximal) whole-cell conductance of *g*_KATP_ = 2.3 nS. In absence of Pinacidil *g*_KATP_ is set equal to 0 nS. (*V*_m_)_pre-stim._ is the membrane voltage in acute slices at equilibrium before drug intervention. Constraints without supporting references were established based on the expected physiological function of each cellular component. The objective function was unweighted by default, except for a scalar weight of 10 assigned to the *V*_m_ residual terms at 10 and 40 mmHg.

As a robustness constraint, model insensitivity to the initial guess was verified at both reference state, the CaPA rest condition (*σ* = 10 mmHg) and the acute-slice baseline (*σ* = 0 mmHg), by re-solving the equilibrium from seven perturbed starting points (initial state varied by *±*5% across fixed seeds) and penalizing any resulting *V*_m_ that deviated from the unperturbed solution by more than 1%, ensuring convergence to a single, seed-independent steady state. The same constraint type was also included during the evaluation of PIN-induced depolarization (using three different seeds). To ensure the model reproduces a physiologically graded (dose-dependent) hyperpolarizing response to K_ATP_ channel activation, rather than an all-or-none response, we imposed a bounding constraint on membrane potential at two submaximal levels of K_ATP_ conductance (*g*_KATP_), corresponding to 33% and 67% of its maximal value. At each submaximal level, the resulting steady-state membrane potential was constrained between the potential under full K_ATP_ activation and the maximum depolarized resting potential (-40 mV). This bounds the response in both directions: a submaximal fraction of active channels cannot hyperpolarize the membrane beyond what full activation achieves, while also requiring an appreciable hyperpolarization relative to rest, preventing the model from collapsing to a flat (non-graded) response across the activation range.

The MLCK and MLCP activity parameters, as well as the vascular wall structural parameters are identified through a second subsequent ‘vascular contractility’ optimization problem, designed to satisfy the PC constriction levels vs pressure (20*→*80 mmHg) under control (CTL) and in absence of extracellular calcium (0Ca) conditions, as reported in [12, 30]. Furthermore, we penalize parameter combinations for which the relative filament sliding *u̅*_fs_ lies outside the physiologically relevant range ([-0.5, 0.1]).

For each optimization problem, the objective function was formed by summing the errors associated with the constraints described above (see Supplementary Information Section 4). The Covariance Matrix Adaptation Evolution Strategy (CMA-ES) is used with multiple (max *≈* 500) iterations to identify the solution minimizing the objective function (as done in our previous work [8]).

We reiterate that only the experimental data reported in Table 1 were used for the model parametrization, while the others (as reported in 3.2) are used for validation.

## 3 Results

### 3.1 Model parametrization

Conductance and *V*_m_ rectification factors for Kir, Kv, TRPC3 and ANO1 channels were firstly identified by using existing whole-cell patch clamp data from rodent cerebral PC [47, 33, 12]. In the absence of cerebral PC-specific data, the remaining model parameters were initialized using data from other cell types, primarily vascular smooth muscle and related expression systems. The cellular signalling and the vascular wall parameters that were identified using the associated optimization problems defined above are reported in the Supplementary Information Section 1. In this section, we present the model parameterization results, demonstrating how well the model captures the fitted experimental data.

#### 3.1.1 Pressure-induced contractility in CaPA

During the parametrization process, model simulations were compared against experimental recordings under the cannulated, pressurized arteriole protocol at two representative intraluminal pressures, 10 mmHg and 40 mmHg and with the addition of GSK1702934A (GSK17). Fig. 2a, upper panel summarizes the steady-state electrophysiological variables under these three conditions. Elevating pressure from 10 to 40 mmHg produced a clear depolarization of the membrane potential in both simulation and experiment (simulated Δ*V*_m_: *≈* 11 mV; experimental Δ*V*_m_: *≈* 13 mV), with the model reproducing the direction and approximate magnitude of the pressure-induced shift, albeit with a more hyperpolarized baseline than observed experimentally in the work by Ferris et al. [12]. This depolarization was accompanied by a rise in intracellular Na^+^ (*≈* 9 to *≈* 15 mM) and a roughly two-fold increase in intracellular Ca^2+^, consistent with pressure-induced activation of depolarizing conductances and secondary Ca^2+^ entry. Intracellular Cl^-^ and SR Ca^2+^ content both decreased modestly over the same pressure range, reflecting redistribution of Cl^-^ and partial store depletion accompanying the elevated cytosolic Ca^2+^ load. With respect to pressure elevation, direct TRPC3 activation by GSK17 (100 nM) caused a similar membrane depolarization, but with an enhanced Na_i_ influx and an alteration of calcium trafficking between the intracellular and store compartments. Decomposition of the total membrane current into its constituent channel and transporter contributions (Fig. 2a, lower panel) demonstrates that the pressure-induced depolarization is dominated by the Na^+^-carrying component of TRPC3 (TRPC3,Na) and by ANO1 (TMEM16A). The inward TRPC3,Na current increased markedly in magnitude with pressure or GSK17 application. Similarly, the Ca^2+^-activated Cl*^−^* current through ANO1 grew with pressure, though ANO1 showed slightly lower activation upon GSK17 treatment. This depolarizing drive was partially offset by an increase in the outward K_Bg_ current and, to a lesser extent, by NaK and PMCA, while Cl_Bg_ current declined toward zero with increasing pressure, consistent with the compensatory relationship between ANO1 and the background Cl^-^ conductance identified during model calibration. The comparison clearly shows that, in our model, TRPC3 provides the primary depolarizing drive during pressure elevation, playing a significantly larger role than TRPM4. As a result of the model parametrization, the simulated steady-state fractional fluorescence change between 20 and 40 mmHg (Δ*F/F*_0_) falls within the experimentally informed range (Fig. 2b). The pharmacological block of voltage-gated Ca^2+^ channels (VGCC) with nifedipine (Nife, 10 *µ*M, modelled as per 3 *µ*M Nimo) abolished the pressure-induced Ca^2+^ signal in the model, driving the simulated steady-state Δ*F/F*_0_ below zero and highlighting the essential role of VGCCs in Ca^2+^ influx. Once the cellular signalling parameters were identified, solving the vascular contractility optimization problem enabled the reconstruction of wall stress/pressure effects on XBs kinetics and capillary diameter deformation (Fig. 2c).

**Figure 2:**
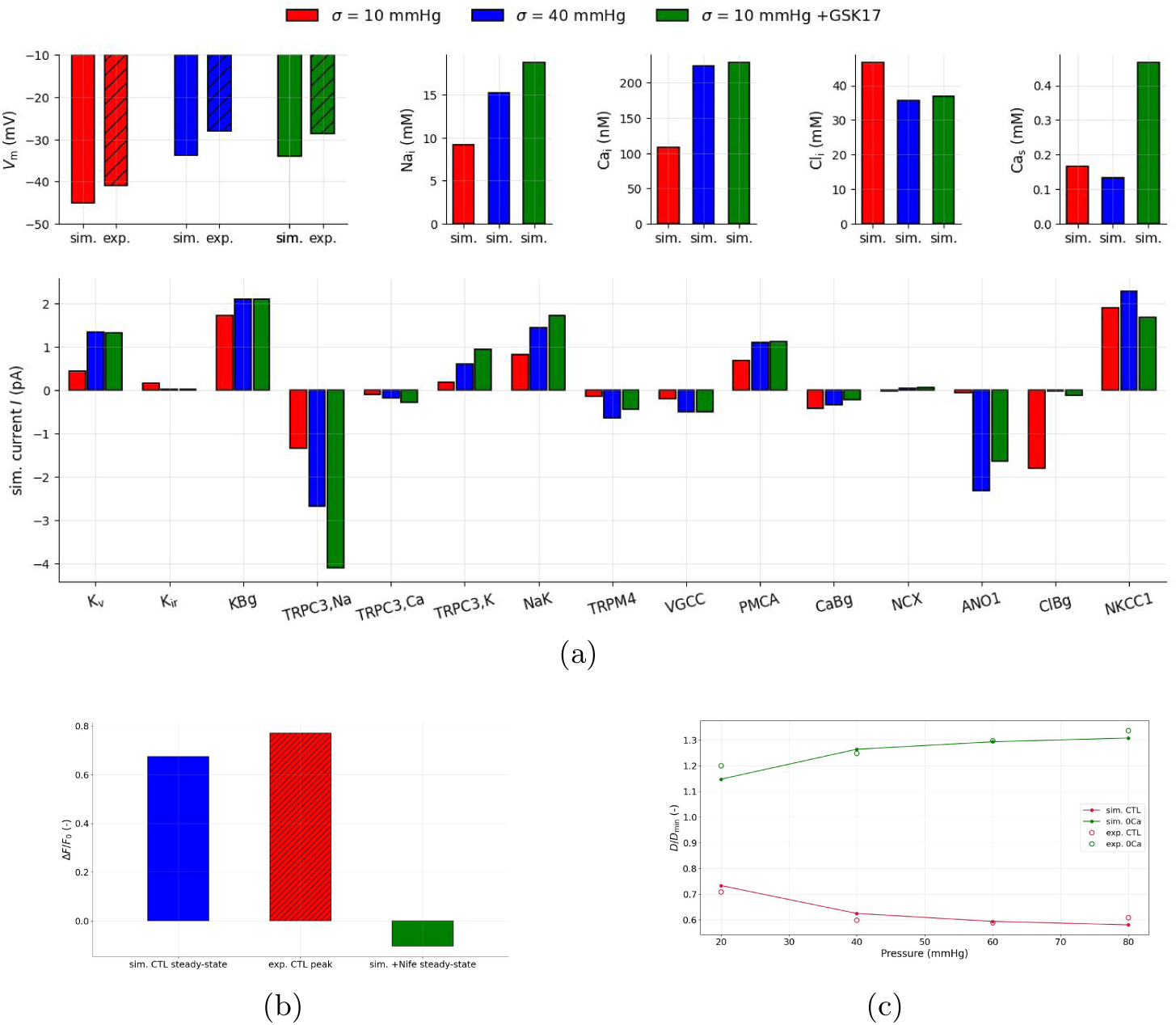
Model parametrization results I: pressure-induced contractility of ensheathing pericytes in the CaPA preparation. **(a)** Electrophysiological balance at different pressure levels and with GSK17. Experimental data (mean values) were obtained from [12]. **(b)** Variation in calcium fluorescence (measured with the GCaMP6f indicator) upon pressure stimulation. The simulated steady-state value is compared vs the experimentally recorded peak from [12], and the simulated effect of VGCC blockage. **(c)** Normalized capillary luminal diameter vs pressure for control (CTL) and extracellular Ca^2+^ removal (0Ca), normalized relative to 5 mmHg values. Experimental data (mean values) are taken from [12] (and re-processed in [30]). The external ion concentrations were set in line with the same study: Na_o_=152.25 mM, K_o_=3 mM, Ca_o_=2 mM, Cl_o_=134 mM. See Table 1 caption for details on modelling drug effects.

#### 3.1.2 Pharmacological modulation in Acute Slices

For model parametrization, we considered changes in calcium signalling induced by 10 nM ET-1 and two associated inhibitors in superfused acute slices (Fig. 3). To replicate this experimental condition, the luminal pressure and wall stress were set to 0 mmHg. We assumed a steady-state ET-1-induced effect, where the ligand concentration is such that the generated IP_3_ concentration (due to solely to the ligand) is twice the SR IP_3_ half-activation constant (*K*_SR,IP3_). While the experimental data (mean values) represent PC responses following approximately 5 minutes of intervention, the model reproduces the steady state intracellular signalling following agonist-stimulation. This temporal mismatch was accounted for in the optimization process by defining a wide acceptance range for fluorescence changes. Application of ET-1 in our model produced a substantial rise in Ca^2+^ fluorescence which falls within the expected experimental range (simulated Δ*F/F*_0_: *≈* 0.42; experimental Δ*F/F*_0_: *≈* 0.61). Co-application of nimodipine, a VGCC blocker, essentially abolished this response in both the model and the experimental data (simulated and experimental Δ*F/F*_0_ both *≈* 0), indicating that the ET-1-evoked Ca^2+^ signal is critically dependent on VGCC-mediated Ca^2+^ entry in both the model and the biological system. In contrast, co-application of the ANO1 (TMEM16A) blocker 10bm (200 nM) reduced but did not eliminate the ET-1 response, confirming an amplifier function for ANO1 in ET-1-evoked Ca^2+^ signalling.

**Figure 3:**
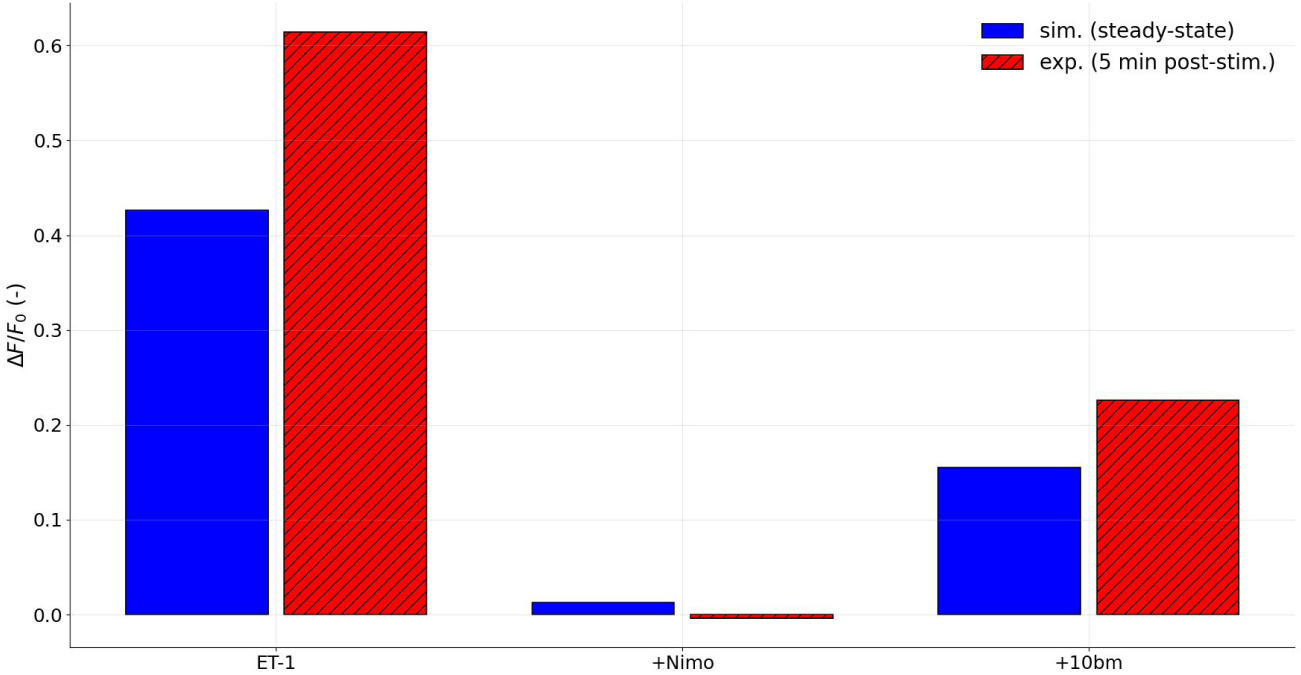
Model parametrization results II: Impact of G_q_CPR activation on pericyte signalling and role of mediators in the acute slices preparation. The experimental data (mean values) were obtained from [32] and represent PC responses following approximately 5 minutes of intervention. Calcium changes were quantified using the fluorescence indicator GCaMP5G. The wall stress in this preparation is assumed to be 0 mmHg. The external ion concentrations are set in line with the same study: Na_o_=151 mM, K_o_=2.5 mM, Ca_o_=2 mM, Cl_o_=132.5 mM. See Table 1 caption for details on modelling drug effects.

### 3.2 Model testing

Model parametrization was carried out by considering specific experimental conditions in CaPA and acute-slice preparations, as reported above. Now we aim to test and validate the current model under new different conditions.

#### 3.2.1 In Vitro Patch-Clamp Conditions

In the model parameter identification, the inactivation and activation properties of the VGCCs *V*_1*/*2VGCC,ina_, *k*_VGCC,ina_, *V*_1*/*2VGCC,act_ and *V_k_*_VGCC,act_ were first initialized by using data from vasa-recta PCs [53]. Given potential discrepancies across tissue types and experimental protocols, the parameters *V*_1*/*2VGCC,ina_, *k*_VGCC,ina_ and *V*_1*/*2VGCC,act_ were re-optimized within admissible bounds derived from earlier studies on PC (or SMC) VGCCs (Fig. 4, upper panel). Parameters optimization led to a shape refinement of these inactivation and activation functions (alongside (*PA*_m_)_VGCC_), which yield a steady state and peak VGCC current which are comparable in shape with previous recordings in cerebral SMCs [45] (Fig. 4, bottom panel). The magnitude ratio between our simulated peak current and the experimentally SMC recording (*≈* 44%) is in line with the ratio reported in a more recent PC-SMC VGCC current comparison (*≈* 51%) [31].

**Figure 4:**
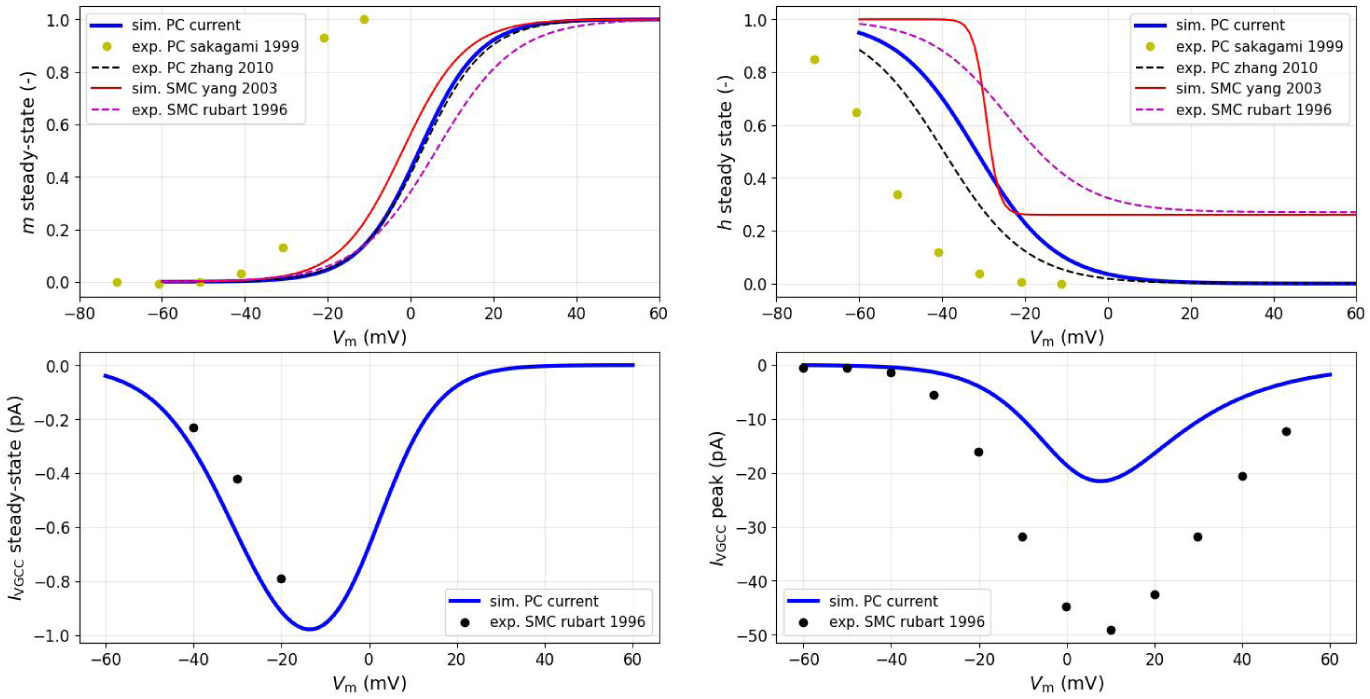
Properties of the modelled VGCCs, with alongside experimental data from different studies on PCs and cerebral SMCs [45, 46, 52, 53]. Top panel: Comparison of voltage-dependent steady-state VGCC activation (right) and inactivation (left) functions between the current model and previous studies. Bottom right panel: Comparison of voltage-dependent steady-state VGCC current between the current model and SMC data from [45]. Bottom left panel: Comparison of voltage-dependent peak VGCC current (*h* = 1) between the current model and SMC data from [45]. Ca_i_ and Ca_o_ were assumed to be 100 nM and 2 mM, respectively.

#### 3.2.2 Ex Vivo Cannulated Vessel Conditions

To evaluate the model’s constriction response beyond the baseline pressure-diameter fit, simulated *D/D*_CTL_ ratios were compared against three independent experimental interventions in cannulated preparations: PIN (pinacidil, a K_ATP_ channel opener), K60/KCl60 (60 mM extracellular K^+^, with and without the accompanying Cl*^−^*), and U46619 (a thromboxane A2-receptor agonist). PIN-induced capillary dilation in CaPA settings was reported in a recent study [27], while Gonzales et al. [17] documented diameter changes induced by KCl60 and U46619 in an en-face retinal microvascular preparation. The maximal K_ATP_ conductance previously derived from in-vitro data [48] and used in the model parametrization may overestimate physiological K_ATP_ currents. Therefore, we also test a scenario where the effective *g*_KATP_ is reduced to 10% of its maximal value (0.23 nS), aligning more closely with the estimates in [27]. To simulate the K60 and KCl60 interventions, we added the corresponding ion concentrations directly to the baseline extracellular bath concentration. Simulating the effect of U46619 on PC contractility is associated with greater uncertainty: rather than resolving the TXA_2_/G_q_/PLC cascade, we approximate its effect as a static elevation of intracellular IP_3_, as done for the endothelin-1 stimulation in Section 3.1.2. U46619 is also known to promote Ca^2+^ sensitization processes via ROCK activation [50], and we modelled a near-maximal contribution to tone by increasing the pressure sensitivity of the ROCK pathway (*K*_MLCP*,σ*_ reduced by 90%). While in CaPA setting the luminal pressure throughout the system is expected to be approximately equal to the inlet pressure (*≈* 40 mmHg), in the retinal preparation (with flow) the wall stress acting on the first- to third-order capillaries remains uncertain. Therefore, in this analysis, diameter changes are evaluated across three pressure levels (20, 30 and 40 mmHg), to reflect the protocol settings in [17, 27]. Our simulations indicate that pressure has a profound impact on the magnitude of the wall response to all three types of interventions (Fig. 5). PIN application caused substantial dilation across the entire pressure range, peaking at 40 mmHg, where baseline control (CTL) diameter was smallest. Intriguingly, a ten-fold reduction in K_ATP_ conductance (*g*_KATP_) slightly increases dilation relative to the reference PIN case, across all pressures. This counterintuitive result reflects that, once VGCC is already fully closed at these hyperpolarized potentials, the dominant residual Ca^2+^ pathway is TRPC3, which is voltage-independent in its gating but driven in magnitude by the transmembrane voltage gradient. A more modest *g*_KATP_ therefore leaves the cell less hyperpolarized, weakening the Ca^2+^ driving force through TRPC3 and reducing myogenic tone generation. Overall, considering both conductances, the predicted dilations range from +38% to +54%, aligning closely with experimental data (+44%). On the other hand, our model predicts greater constriction when K60/KCl60 and U46619 are applied to lower vessel pressures, suggesting a potential limit on steady-state calcium-dependent contractile capacity. Nevertheless, the predicted diameter reductions induced by adding K60/KCl60 align closely with the experimental recordings, particularly for *σ* between 20 and 30 mmHg. Without ROCK modulation, our model predicts U46619-induced constrictions between -4 % and -12 % across the pressure range, which appear severely blunted compared to the experimental data (-34%). Adding a U46619induced ROCK-modulation effect (in which *K*_MLCP*,σ*_ is decreased ten-fold to represent a maximal enhancement of ROCK-mediated Ca^2+^-sensitization), substantially diminishes the gap between simulated and experimental results.

**Figure 5:**
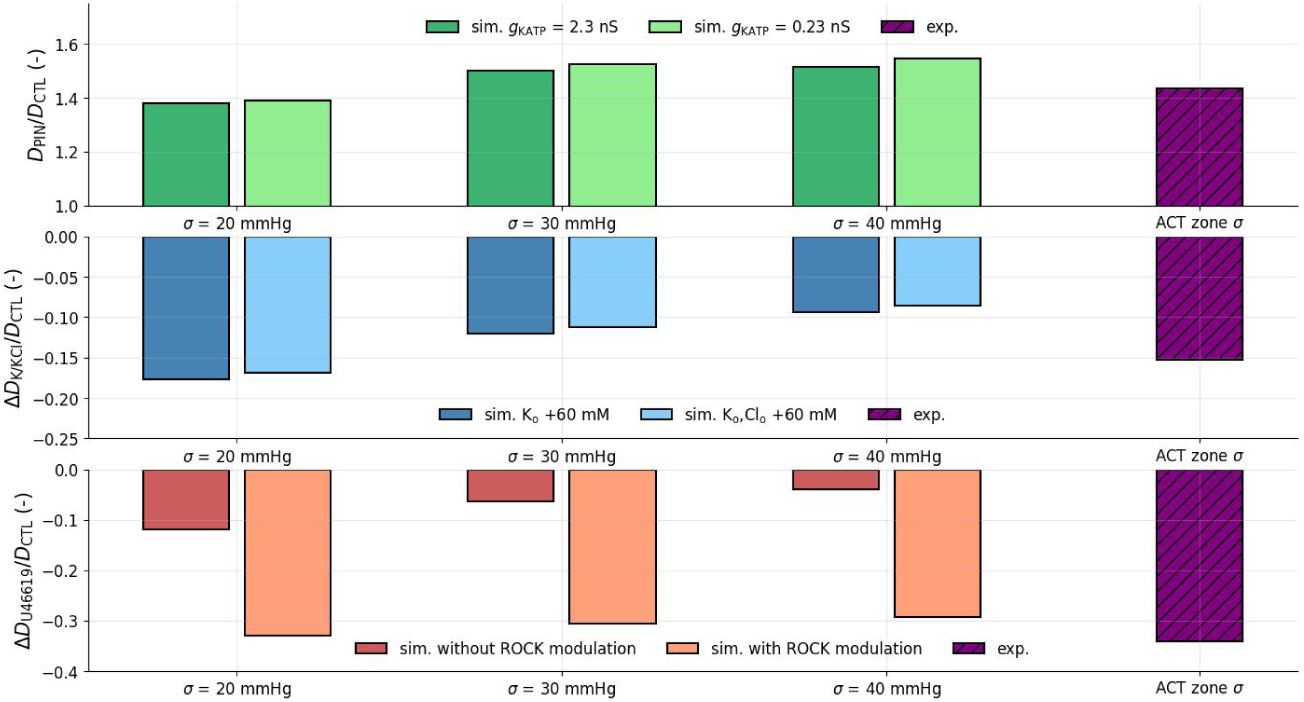
Impact of PIN, K60/KCl60 and U46619 on PC contractility and luminal diameter. Experimental data (mean values) for PIN-induced dilation were obtained from CaPA settings [27]. The experimental data (mean values) for K60/KCl60and U46619-induced constriction were obtained from en-face retinal microvascular preparation. For the PIN-induced dilation, the external ion concentrations matched those used to assess pressure effect in the CaPA preparation (see Table 1 caption), as described in [26, 12]. For the other two interventions, the external ion concentrations were set in line with [17]: Na_o_=151 mM, K_o_ = 2.5 mM, Ca_o_=1.8 mM, Cl_o_ = 130.6 mM.

#### 3.2.3 In Vivo Conditions

Here we predict the in vivo effect of systemic VGCC blockade (nimodipine, 3 *µ*M) on PC intracellular calcium signalling and diameter regulation. To account for the expected relaxation of the upstream vasculature, VGCC blockade was simulated across a range of physiological baseline pressures (*σ*_pre-stim._ = 20, 25, and 30 mmHg) paired with three superimposed post-stimulation pressure step elevations (Δ*σ*_post-stim._ = 0, 3, and 5 mmHg). Results were validated against in vivo imaging measurements (Fig. 6). Simulating nimodipine application produced a consistent, negative shift in simulated Δ*F/F*_0_, the magnitude of which increased at higher baseline luminal pressures. The potential effect of upstream vascular relaxation (Δ*σ*_post-stim._= 3, 5 mmHg) attenuated the nimodipine-induced Ca^2+^ variation across all baseline pressure conditions.

**Figure 6:**
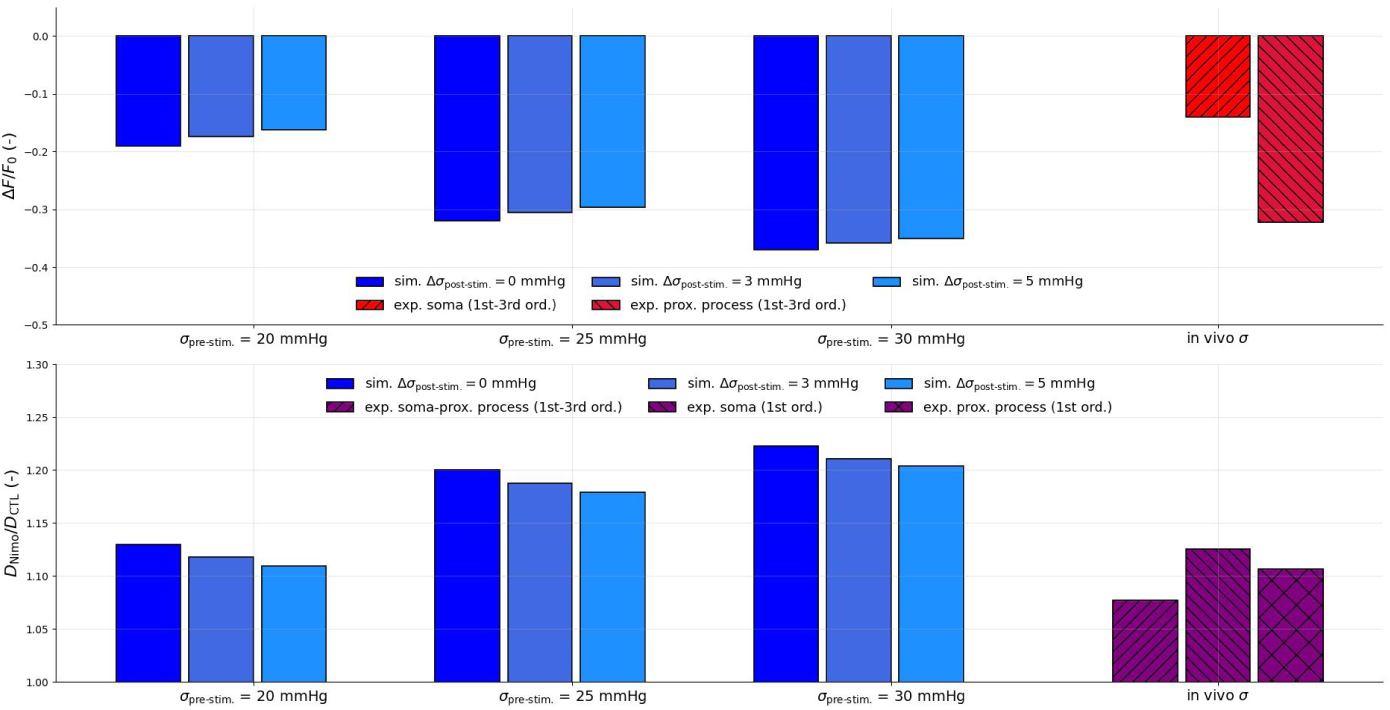
Impact of VGCC blockage on change in calcium levels and luminal diameter under in vivo conditions. Calcium changes were quantified using the fluorescence indicator GCaMP5G. Experimental data (mean values) were obtained from [32]. External ion concentrations are assumed to match those used in the CaPA preparation [26, 12]: Na_o_=152.25 mM, K_o_=3 mM, Ca_o_=2 mM, Cl_o_=134 mM. See Table 1 caption for details on modelling Nimo effect on PC signalling. *σ*_pre-stim._ : *σ* before Nimo application; Δ*σ*_post-stim._ : *σ* variation after Nimo application.

The simulated values for *σ*_pre-stim._ = 20 and 25 mmHg bracket the two in vivo experimental estimates, which showed a smaller reduction in the soma (*≈ −*14%) and a larger reduction in processes (*≈ −*32%), whereas the higher pressure case markedly overestimated the decrease in Ca^2+^ levels. The corresponding diameter response, expressed as the ratio of vessel diameter under VGCC block to control diameter (*D*_Nimo_*/D*_CTL_), showed a consistent dilation upon VGCC block across all simulated conditions (ratios ranging from *≈* 11 % to *≈* 22 %), with the largest dilation predicted at the highest baseline pressure and smallest Δ*σ*_post-stim._, and progressively smaller dilation at lower baseline pressure and larger Δ*σ*_post-stim._. While experimental in vivo diameter reductions (*≈* 10%) were slightly smaller than most simulated values, all cases remained of comparable magnitude, with the *σ*_pre-stim._ = 20 mmHg condition showing the closest agreement. For consistency, external ion concentrations were aligned with the CaPA protocol to faithfully reflect physiological extracellular levels; however, non-negligible variations in bath composition may significantly alter system behaviour. Taken together, these results indicate that the model can reliably predict the direction and approximate magnitude of VGCC-dependent vasodilation observed in vivo, highlighting the role of wall stress (here approximated by pressure) in the pharmacological modulation of microvascular tone. Recent in vivo experiments [51] suggest that ANO1 channels play a key role in basal tone in mid-capillary pericytes. We therefore also computationally probed, with the current model parametrization, the impact of ANO1 channel inhibition (via 10bm) on PC capillary diameter. Under these conditions, we recorded a modest relaxation across the same pressure field considered above, with a maximum dilation of 5.8 % at 25 mmHg. This could suggest that, under physiological conditions, the contribution of ANO1 may be more limited in upstream PCs, and that its loss may be compensated for by other ionic mechanisms. However, the current model parametrization did not directly account for experimental data on the pressure-dependent contribution of Cl*^−^*. Therefore, additional experimental data on ensheathing pericytes are needed to guide further investigation.

## 4 Discussion

Advancing our understanding of cerebral microvascular flow regulation and its impairment requires developing robust theoretical models to integrate diverse types of data across different experimental conditions. The sparsity and uncertainty associated with experimental data represent a major obstacle regarding model structure and parameter identifiability. On top of that, the current contractility paradigm for mural cells relies on positive feedback mechanisms to significantly elevate intracellular calcium and facilitate tone generation. From a physiological standpoint, these cells possess classic features of ‘robust’ control systems, in which small perturbations in extracellular concentrations and channel gating properties produce gradual, adaptive responses. However, the presence of positive feedback loops necessary to elevate intracellular calcium concentration and activate the contractile machinery, alongside non-linear membrane currents, introduces the potential for system bistability [10], posing a challenge to overall model robustness.

In this study, we introduced a new multiscale computational framework for quantifying PC contractility by devising an optimization strategy that accounts for diverse experimental data and physiological constraints. The model was successfully validated across four distinct experimental scenarios under both ex vivo and in vivo conditions using different pharmacological interventions, demonstrating close agreement with recorded data. Although our model incorporates several components from previous studies [52, 28, 8], its architecture includes novel elements, such as localized calcium activation of ANO1 and GPCR-biased signaling, which could be refined in future work as more detailed PC characterization becomes available. Compared to earlier modeling studies on microvascular contractility [40, 30], this work provides a new perspective on how different pharmacological agents (such as pinacidil, elevated external K^+^, U46619, and nimodipine) impact cell signaling and contractility alongside mechanical stress. The demonstrated predictive capabilities, together with the calibrated behavior, support the model’s generalizability. Our simulation results indicate that PC mechanical activation via wall stress must be factored in when quantitative investigations of vasoactive compounds are pursued. The slightly blunted pressure-induced depolarization yielded by our model parametrization suggests that the description of the underlying currents may still be incomplete. In this context, additional data characterizing other contributors to pressureor flow-induced depolarization as well as calcium-desensitization mechanisms are necessary to advance the model’s predictive capabilities. From a physiological perspective, the model could also be refined to distinguish between specific VGCC subtypes (i.e., L-type vs T-type), while ligand-mediated GPCR activation deserves a more detailed description to sufficiently capture drug-specific kinetics. Furthermore, ensheathing pericytes were assumed here to fully wrap the endothelial tube as an infinitesimally long ring, without accounting for the attenuation of pinching forces toward their processes. Future modeling efforts should incorporate more realistic PC mechanics and geometry [11] to assess their impact on microvessel deformation and flow regulation.

As noted earlier, describing the processes underlying vascular control is challenged by the multiscale complexity and intrinsic uncertainty of the system itself. Data scarcity often leads to parameter identifiability issues, while positive feedback control leading to potential bistable behavior can severely undermine the model’s utility. We addressed these challenges in part by defining a parameter optimization strategy that incorporated diverse experimental data and guaranteed a baseline level of robustness under variable initial conditions. By imposing numerous physiological constraints during parameterization, we avoided anchoring key channel parameters (such as channel conductances) to in vitro whole-cell measurements, which may reflect non-physiological properties. The stochastic nature of the optimizer enabled the exploration of different parameter combinations, but it can also identify multiple parameter sets as independently valid solutions. The primary objective of this study was to demonstrate that the proposed optimization strategy could yield at least one parameter set capable of capturing distinct electrophysiological and vascular tissue data with reasonable accuracy; comprehensive parameter exploration was beyond the scope of this work. Nevertheless, sensitivity analysis of the parameter space and thorough evaluation of the bistable regime are certainly warranted. The next natural step in model refinement and testing will involve evaluating pharmacomechanical stimuli within realistic microvascular networks, establishing a foundation for translational studies. Indeed, the proposed framework provides a basis for designing therapies to mitigate tissue ischemia associated with a range of cerebrovascular pathologies, such as stroke and vascular dementia [33, 32].

