## Supplementary Information for "Predicting Cerebral Pericyte Contractility Across Experimental and Physiological Conditions: an in-silico framework"

### Supplementary Information: Model Parameters, Governing Equations and Objective Function Definition

#### S1 Fitted Model Parameters

Table S1: Fitted model parameters identified through the cellular signalling optimization problem. Bounds are expressed as multiplicative factors on the baseline (reference) value; the identified value is baseline × identified factor. The best objective residual is 0.0138.

| # | Parameter | Description | Lower bound | Upper bound | Identified factor | Baseline | Units | Identified value |
|---|---|---|---|---|---|---|---|---|
| 1 | $V_{1/2,\mathrm{VGCC,act}}$ | VGCC activation half-activation voltage | 0.700 | 1.250 | 0.7608 | 2.8 | mV | 2.130 |
| 2 | $(PA_\mathrm{m})_\mathrm{VGCC}$ | VGCC membrane permeability-area product | 0.250 | 1.500 | 0.7518 | $1.5 \times 10^{-10}$ | $\mathrm{cm^3/s}$ | $1.13 \times 10^{-10}$ |
| 3 | $V_{1/2,\mathrm{VGCC,ina}}$ | VGCC inactivation half-inactivation voltage | 0.800 | 1.250 | 0.8008 | $-39.6$ | mV | $-31.71$ |
| 4 | $k_\mathrm{VGCC,ina}$ | VGCC inactivation slope factor | 0.600 | 1.250 | 0.9652 | $-10$ | mV | $-9.652$ |
| 5 | $g_\mathrm{KBg}$ | Background $K^+$ conductance | 0.010 | 2.000 | 0.3318 | 0.1 | nS | $3.32 \times 10^{-2}$ |
| 6 | $g_\mathrm{Kir}$ | Kir channel conductance | 0.500 | 1.000 | 0.8940 | 0.426 | $\mathrm{nS/mM^{0.5}}$ | 0.3808 |
| 7 | $k_\mathrm{Kir}$ | Kir activation slope factor | 0.750 | 1.500 | 0.9830 | $-5.25$ | mV | $-5.161$ |
| 8 | $(PA_\mathrm{m})_\mathrm{ANO1}$ | ANO1 membrane permeability-area product | 0.010 | 10.000 | 5.5188 | $3.9 \times 10^{-9}$ | $\mathrm{cm^3/s}$ | $2.15 \times 10^{-8}$ |
| 9 | $K_\mathrm{ANO1,Ca}$ | ANO1 $Ca^{2+}$ half-activation constant | 0.500 | 1.500 | 1.1285 | $5.0 \times 10^{-4}$ | mM | $5.64 \times 10^{-4}$ |
| 10 | $(PA_\mathrm{m})_\mathrm{ClBg}$ | Background $Cl^-$ permeability-area product | 0.010 | 1.400 | 1.2987 | $2.8 \times 10^{-13}$ | $\mathrm{cm^3/s}$ | $3.64 \times 10^{-13}$ |
| 11 | $K_{\mathrm{G}\sigma}$ | $G_q$ stress half-activation constant | 1.000 | 5.000 | 3.9991 | 10 | mmHg | 39.99 |
| 12 | $(PA_\mathrm{m})_\mathrm{TRPC3}$ | TRPC3 membrane permeability-area product | 0.250 | 1.250 | 0.3540 | $5.0 \times 10^{-13}$ | $\mathrm{cm^3/s}$ | $1.77 \times 10^{-13}$ |
| 13 | $\alpha_\mathrm{TRPC3}$ | TRPC3 constitutive baseline activity | 0.200 | 2.000 | 1.7145 | 0.1 | (–) | 0.1714 |
| 14 | $n_{\mathrm{G}\sigma}$ | $G_q$ stress activation Hill coefficient | 0.500 | 2.000 | 0.8468 | 2 | (–) | 1.694 |
| 15 | $\alpha_\mathrm{TRPM4}$ | TRPM4 constitutive baseline activity | 0.100 | 1.300 | 1.1448 | 0.3 | (–) | 0.3434 |
| 16 | $(PA_\mathrm{m})_\mathrm{TRPM4}$ | TRPM4 membrane permeability-area product | 0.100 | 5.000 | 1.8346 | $1.0 \times 10^{-13}$ | $\mathrm{cm^3/s}$ | $1.83 \times 10^{-13}$ |
| 17 | $n_{\mathrm{TRPM4},\sigma}$ | TRPM4 stress activation Hill coefficient | 0.500 | 2.000 | 0.6626 | 2 | (–) | 1.325 |
| 18 | $J_\mathrm{IP3R,max}$ | Maximal $IP_3R$ $Ca^{2+}$ release flux rate | 0.500 | 5.000 | 3.6396 | 0.25 | mM/ms | 0.9099 |
| 19 | $\mathrm{IP}_{3,\mathrm{G}\sigma,\mathrm{max}}$ | Maximal stress-induced $IP_3$ production | 0.100 | 1.500 | 0.3170 | $3.0 \times 10^{-4}$ | mM | $9.51 \times 10^{-5}$ |
| 20 | $K_\mathrm{SERCA}$ | SERCA pump $Ca^{2+}$ half-activation constant | 0.500 | 2.000 | 0.5984 | $4.0 \times 10^{-4}$ | mM | $2.39 \times 10^{-4}$ |
| 21 | $L_\mathrm{SR}$ | SR $Ca^{2+}$ leak rate constant | 0.100 | 2.000 | 0.7362 | $6.0 \times 10^{-5}$ | 1/ms | $4.42 \times 10^{-5}$ |
| 22 | $\beta_\mathrm{ANO1}$ | ANO1 local concentration scaling factor | 0.050 | 2.000 | 0.0593 | 100 | (–) | 5.928 |
| 23 | $K_\mathrm{Orai}$ | Orai store $Ca^{2+}$ half-inactivation constant | 0.100 | 5.000 | 1.6990 | 0.25 | mM | 0.4247 |
| 24 | $(PA_\mathrm{m})_\mathrm{CaBg}$ | Background $Ca^{2+}$ permeability-area product | 0.200 | 2.000 | 0.5736 | $7.1 \times 10^{-13}$ | $\mathrm{cm^3/s}$ | $4.07 \times 10^{-13}$ |
| 25 | $K_\mathrm{NaK,Na}$ | $Na^+/K^+$-ATPase $Na^+$ half-activation constant | 0.500 | 2.500 | 2.0675 | 10 | mM | 20.68 |
| 26 | $g_\mathrm{NCX}$ | $Na^+/Ca^{2+}$ exchanger activity constant | 0.050 | 10.000 | 0.0587 | $1.0 \times 10^{-3}$ | (–) | $5.87 \times 10^{-5}$ |
| 27 | $L_\mathrm{NKCC1}$ | NKCC1 cotransporter maximal transport rate | 0.250 | 2.000 | 1.7778 | 2.5 | mM/ms | 4.444 |
| 28 | $K_\mathrm{NKCC1,ina}$ | NKCC1 intracellular $Cl^-$ inactivation constant | 0.800 | 1.250 | 1.2215 | 25 | mM | 30.54 |
| 29 | $I_\mathrm{PMCA,max}$ | PMCA pump maximal current | 0.333 | 1.000 | 0.4305 | 6 | pA | 2.583 |
| 30 | $K_\mathrm{PMCA}$ | PMCA pump $Ca^{2+}$ half-activation constant | 0.500 | 1.500 | 1.4972 | $2.0 \times 10^{-4}$ | mM | $2.99 \times 10^{-4}$ |

Table S2: Fitted model parameters identified through the vascular contractility optimization problem. Bounds are expressed as multiplicative factors on the baseline (reference) value; the identified value is baseline × identified factor. Best objective residual is 0.0011973. $\kappa_{ps}$ is set equal to 3.8 $\times 10^6$ dyne/cm$^2$.

| # | Parameter | Description | Lower bound | Upper bound | Identified factor | Baseline | Units | Identified value |
|---|---|---|---|---|---|---|---|---|
| 0 | $R_o$ | Reference outer radius | 0.900 | 2.000 | 1.7396 | 3.0 | $\mu$m | 5.219 |
| 1 | $h_w$ | Reference wall thickness-radius ratio | 0.250 | 2.000 | 0.4703 | 0.25 | (–) | 0.1176 |
| 2 | $c_0$ | Passive stiffness coefficient | 0.050 | 6.000 | 2.3823 | $9.1633 \times 10^4$ | dyne/cm$^2$ | $2.18 \times 10^5$ |
| 3 | $c_1$ | Passive exponential parameter | 0.250 | 12.000 | 0.2503 | $3.15 \times 10^4$ | dyne/cm$^2$ | $7.88 \times 10^3$ |
| 4 | $c_2$ | Passive exponent scaling factor | 0.010 | 10.000 | 5.6979 | 1.292 | (–) | 7.362 |
| 5 | $K_{MLCK,Ca}$ | MLCK $Ca^{2+}$ half-activation constant | 0.500 | 2.500 | 2.4994 | $2.0 \times 10^{-4}$ | mM | $5.00 \times 10^{-4}$ |
| 6 | $n_{MLCK}$ | MLCK Hill coefficient | 0.500 | 2.000 | 0.7651 | 2 | (–) | 1.530 |
| 7 | $K_{MLCP,\sigma}$ | MLCP stress half-activation constant | 0.500 | 2.000 | 1.4969 | 40 | mmHg | 59.88 |
| 8 | $n_{MLCP}$ | MLCP Hill coefficient | 0.500 | 2.000 | 0.5008 | 2 | (–) | 1.002 |
| 9 | $\alpha_{MLCP}$ | MLCP baseline activity level | 0.250 | 2.000 | 0.7894 | 0.5 | (–) | 0.3947 |
| 10 | $k_t/\kappa_{ps}$ | XB stiffness-power stroke ratio | 0.500 | 20.000 | 19.6683 | 1.0 | (–) | 19.67 |

#### S2 Fixed Model Parameters

Table S3: Cell geometry, baseline ionic concentrations and fluorescence indicator parameters.

| Parameter | Description | Value | Units |
|---|---|---|---|
| $C_m$ | Membrane capacitance | 10.1 | pF |
| $v_i$ | Cytosolic volume | 1.0 | pL |
| $v_{Cai}$ | Effective $Ca^{2+}$ cytosolic volume | 0.7 | pL |
| $v_{SR}$ | SR volume | 0.07 | pL |
| $\beta_{buff}$ | $Ca^{2+}$ buffering factor | 1/80 | – |
| $V_m^{ref}$ | Initial membrane potential | 140 | mM |
| $Na_i^{bas}$ | Initial intracellular $Na^+$ | 8 | mM |
| $K_i^{bas}$ | Initial intracellular $K^+$ | 140 | mM |
| $Cl_i^{bas}$ | Initial intracellular $Cl^-$ | 40 | mM |
| $Ca_i^{bas}$ | Initial intracellular $Ca^{2+}$ | $10^{-4}$ | mM |
| $Ca_s^{bas}$ | Initial SR $Ca^{2+}$ | 0.5 | mM |
| $R_f$ | Fluorescence scaling factor | 10 | – |
| $K_{df}$ | Indicator half-saturation constant | 375 (GCaMP6f), 447 (GCaMP5G) | nM |
| $n_f$ | Indicator Hill coefficient | 2.27 (GCaMP6f), 2.46 (GCaMP5G) | – |

Table S4: Parameters for ion channels, transporters and exchangers.

| Parameter | Description | Value | Units |
|---|---|---|---|
| $V_{1/2,\mathrm{Kv,act}}$ | Kv half-activation voltage | 15.37 | mV |
| $k_{\mathrm{Kv,act}}$ | Kv activation slope factor | 12.42 | mV |
| $g_{\mathrm{Kv}}$ | Kv conductance | 1.127 | nS |
| $n_{\mathrm{Kv}}$ | Kv activation exponent | 1 | – |
| $g_{\mathrm{Kir}}$ | Kir conductance | 0.426 | nS |
| $\Delta V_{\mathrm{Kir}}$ | Kir offset voltage | 25 | mV |
| $n_{\mathrm{Kir}}$ | Kir $K_{\mathrm{o}}$ dependence | 0.5 | – |
| $k_{\mathrm{VGCC,act}}$ | VGCC activation slope factor | 7.3 | mV |
| $\gamma_{\mathrm{TRPC3},j}$ ($j \in \{\mathrm{Na, Ca, K}\}$) | TRPC3 relative permeability factor | 1.0 (Na, K), 1.6 (Ca) | – |
| $K_{\mathrm{TRPM4},\sigma}$ | TRPM4 stress half-activation constant | 20 | mmHg |
| $K_{\mathrm{TRPM4,Ca}}$ | TRPM4 $Ca^{2+}$ half-activation constant | $5 \times 10^{-4}$ | mM |
| $n_{\mathrm{TRPM4,Ca}}$ | TRPM4 $Ca^{2+}$ Hill coefficient | 2 | – |
| $V_{1/2,\mathrm{CaBg}}$ | CaBg half-rectification voltage | -70 | mV |
| $k_{\mathrm{CaBg}}$ | CaBg rectification slope factor | 2.5 | mV |
| $n_{\mathrm{Orai}}$ | Orai Hill coefficient | 1 | – |
| $V_{1/2,\mathrm{ANO1}}$ | ANO1 half-rectification voltage | 0.0 | mV |
| $k_{\mathrm{ANO1}}$ | ANO1 rectification slope factor | 41.84 | mV |
| $n_{\mathrm{ANO1,Ca}}$ | ANO1 $Ca^{2+}$ Hill coefficient | 8 | – |
| $n_{\mathrm{PMCA,Ca}}$ | PMCA pump $Ca^{2+}$ Hill coefficient | 1 | – |
| $I_{\mathrm{NaK,max}}$ | Maximum NaK pump current | 8.0 | pA |
| $K_{\mathrm{NaK,Ko}}$ | NaK pump $K^{+}$ affinity | 1.5 | mM |
| $n_{\mathrm{NaK,Ko}}$ | NaK pump $K^{+}$ Hill coefficient | 1.0 | – |
| $n_{\mathrm{NaK,Na}}$ | NaK pump $Na^{+}$ affinity | 1.5 | mM |
| $\gamma_{\mathrm{NCX}}$ | NCX voltage-dependence partition coefficient | 0.45 | – |
| $d_{\mathrm{NCX}}$ | NCX saturation constant | $3 \times 10^{-4}$ | $\mathrm{mM}^{-4}$ |
| $I^{max}_{\mathrm{SERCA,max}}$ | SERCA pump maximum uptake | 6.0 | pA |
| $n_{\mathrm{SERCA}}$ | SERCA pump $Ca^{2+}$ Hill coefficient | 2 | – |
| $K_{\mathrm{SR,IP3}}$ | SR $IP_3$ half-activation constant | $1.2 \times 10^{-4}$ | mM |
| $K_{\mathrm{SR,Ca,act}}$ | SR $Ca^{2+}$ half-activation constant | $1.7 \times 10^{-4}$ | mM |
| $K_{\mathrm{SR,Ca,ina}}$ | SR $Ca^{2+}$ Half-inactivation constant | $1.0 \times 10^{-4}$ | mM |

### S3 Governing Equations

The Hill and Boltzmann sigmoid activation functions $H(x; n, K)$ and $B(V_{\mathrm{m}}; V_{1/2}, k)$ are defined once and reused throughout

$$H(x; n, K) = \frac{x^n}{x^n + K^n}, \qquad B(V_{\mathrm{m}}; V_{1/2}, k) = \frac{1}{1 + e^{\left(\frac{V_{1/2} - V_{\mathrm{m}}}{k}\right)}}. \tag{S1}$$

In this study, the steady-state value of each activation and inactivation gating variable is modeled using a Boltzmann function under the quasi-steady-state approximation.

#### S3.1 Membrane potential dynamics

The model describes the dynamics of membrane potential ($V_{\mathrm{m}}$), intracellular sodium ($\mathrm{Na_i}$), calcium ($\mathrm{Ca_i}$), chloride ($\mathrm{Cl_i}$), potassium ($\mathrm{K_i}$), and sarcoplasmic reticulum calcium ($\mathrm{Ca_s}$). Membrane potential follows the Hodgkin–Huxley formalism,

$$C_{\mathrm{m}} \frac{dV_{\mathrm{m}}}{dt} = -I_{\mathrm{tot}}, \tag{S2}$$

where

$$I_{\mathrm{tot}} = I_{\mathrm{K}} + I_{\mathrm{Na}} + I_{\mathrm{Ca}} + I_{\mathrm{Cl}}. \tag{S3}$$

The lumped membrane currents are defined as

$$I_{\mathrm{K}} = I_{\mathrm{Kv}} + I_{\mathrm{Kir}} + I_{\mathrm{KBg}} + I_{\mathrm{KATP}} + I_{\mathrm{TRPC3,K}} - 2I_{\mathrm{NaK}} - \tfrac{1}{2}I_{\mathrm{NKCC1}}, \tag{S4}$$

$$I_{\mathrm{Na}} = I_{\mathrm{TRPC3,Na}} + 3I_{\mathrm{NaK}} + 3I_{\mathrm{NCX}} + I_{\mathrm{TRPM4}} - \tfrac{1}{2}I_{\mathrm{NKCC1}}, \tag{S5}$$

$$I_{\mathrm{Ca}} = I_{\mathrm{TRPC3,Ca}} + I_{\mathrm{VGCC}} + I_{\mathrm{PMCA}} + I_{\mathrm{CaBg}} - 2I_{\mathrm{NCX}}, \tag{S6}$$

$$I_{\mathrm{Cl}} = I_{\mathrm{ANO1}} + I_{\mathrm{ClBg}} + I_{\mathrm{NKCC1}}. \tag{S7}$$

#### S3.2 Intracellular ion dynamics

The ion concentrations are governed by the mass-balance equations

$$\frac{d\mathrm{Na_i}}{dt} = -\frac{I_{\mathrm{Na}}}{Fv_{\mathrm{i}}}, \tag{S8}$$

$$\frac{d\mathrm{Ca_i}}{dt} = -\beta_{\mathrm{buff}}\,\frac{I_{\mathrm{Ca}} - I_{\mathrm{SR}}}{z_{\mathrm{Ca}}Fv_{\mathrm{Cai}}}, \tag{S9}$$

$$\frac{d\mathrm{Cl_i}}{dt} = -\frac{I_{\mathrm{Cl}}}{z_{\mathrm{Cl}}Fv_{\mathrm{i}}}, \tag{S10}$$

$$\frac{d\mathrm{K_i}}{dt} = -\frac{I_{\mathrm{K}}}{Fv_{\mathrm{i}}}, \tag{S11}$$

$$\frac{d\mathrm{Ca_s}}{dt} = -\frac{I_{\mathrm{SR}}}{z_{\mathrm{Ca}}Fv_{\mathrm{SR}}}, \tag{S12}$$

where $I_{\mathrm{SR}} = (J_{\mathrm{IP_3R}} + J_{\mathrm{SRL}})z_{\mathrm{Ca}}Fv_{\mathrm{Cai}} - I_{\mathrm{SERCA}}$.

#### S3.3 Membrane currents

The Ohmic (linear) background and voltage-gated currents are

$$I_{\mathrm{Kv}} = g_{\mathrm{Kv}}\, m_{\mathrm{Kv}}^{n_{\mathrm{Kv}}}\,(V_{\mathrm{m}} - E_{\mathrm{K}}), \tag{S13}$$

$$I_{\mathrm{Kir}} = g_{\mathrm{Kir}}\, \mathrm{K_o}^{n_{\mathrm{Kir}}}\, B(V_{\mathrm{m}}; V_{1/2,\mathrm{Kir}}, k_{\mathrm{Kir}})\,(V_{\mathrm{m}} - E_{\mathrm{K}}), \tag{S14}$$

$$I_{\mathrm{KATP}} = g_{\mathrm{KATP}}\,(V_{\mathrm{m}} - E_{\mathrm{K}}), \tag{S15}$$

$$I_{\mathrm{KBg}} = g_{\mathrm{KBg}}\,(V_{\mathrm{m}} - E_{\mathrm{K}}), \tag{S16}$$

where $m_{\mathrm{Kv}}$ is the Kv activation gating variable, $B(V_{\mathrm{m}}; V_{1/2,\mathrm{Kir}}, k_{\mathrm{Kir}})$ is the voltage-dependent rectification of $\mathrm{K_{ir}}$ channel, with $V_{1/2,\mathrm{Kir}} = \Delta V_{\mathrm{Kir}} + E_{\mathrm{K}}$.

Ion currents through GHK-type permeation pathways are written using a single Goldman–Hodgkin–Katz current relation,

$$\mathrm{GHK}(V_{\mathrm{m}}, z, C_i, C_o) = z^2\,\frac{V_{\mathrm{m}}F^2}{RT}\,\frac{C_i - C_o\exp(-zFV_{\mathrm{m}}/RT)}{1 - \exp(-zFV_{\mathrm{m}}/RT)}, \tag{S17}$$

so that

$$I_{\mathrm{TRPC3},j} = (PA_{\mathrm{m}})_{\mathrm{TRPC3}}\,\gamma_{\mathrm{TRPC3},j}\,f_{\mathrm{TRPC3},\sigma}(\sigma)\,\mathrm{GHK}\,(z_j, C_{i,j}, C_{o,j},,) \qquad j \in \{\mathrm{Na},\mathrm{Ca},\mathrm{K}\}, \tag{S18}$$

$$I_{\mathrm{VGCC}} = (PA_{\mathrm{m}})_{\mathrm{VGCC}}\,m_{\mathrm{VGCC}}\,h_{\mathrm{VGCC}}\,\mathrm{GHK}\,(z_{\mathrm{Ca}}, \mathrm{Ca_i}, \mathrm{Ca_o},,) \tag{S19}$$

$$I_{\mathrm{CaBg}} = (PA_{\mathrm{m}})_{\mathrm{CaBg}}\,H(\mathrm{Ca_s}; n_{\mathrm{Orai}}, K_{\mathrm{Orai}})\,B(V_{\mathrm{m}}; V_{1/2,\mathrm{CaBg}}, k_{\mathrm{CaBg}})\,\mathrm{GHK}\,(z_{\mathrm{Ca}}, \mathrm{Ca_i}, \mathrm{Ca_o},,) \tag{S20}$$

$$I_{\mathrm{ANO1}} = (PA_{\mathrm{m}})_{\mathrm{ANO1}}\,f_{\mathrm{ANO1,Ca}}\,B(V_{\mathrm{m}}; V_{1/2,\mathrm{ANO1}}, k_{\mathrm{ANO1}})\,\mathrm{GHK}\,(z_{\mathrm{Cl}}, \mathrm{Cl_i}, \mathrm{Cl_o},,) \tag{S21}$$

$$I_{\mathrm{ClBg}} = (PA_{\mathrm{m}})_{\mathrm{ClBg}}\,\mathrm{GHK}\,(z_{\mathrm{Cl}}, \mathrm{Cl_i}, \mathrm{Cl_o},,) \tag{S22}$$

$$\begin{aligned} I_{\mathrm{TRPM4}} = {} & (PA_{\mathrm{m}})_{\mathrm{TRPM4}}\,H(\mathrm{Ca_i}; n_{\mathrm{TRPM4,Ca}}, K_{\mathrm{TRPM4,Ca}}) \\ & \cdot H(\sigma; n_{\mathrm{TRPM4},\sigma}, K_{\mathrm{TRPM4},\sigma})\,\mathrm{GHK}\,(z_{\mathrm{Na}}, \mathrm{Na_i}, \mathrm{Na_o}, .) \end{aligned} \tag{S23}$$

where $m_{\mathrm{VGCC}}$ and $h_{\mathrm{VGCC}}$ are the VGCC activation and inactivation gating variables.

The SR currents are defined as follows:

$$J_{\mathrm{SRL}} = L_{\mathrm{SR}}\,(\mathrm{Ca_s} - \mathrm{Ca_i}), \tag{S24}$$

$$P_{\mathrm{IP3R}} = \frac{K_{\mathrm{SR,Ca,ina}}}{K_{\mathrm{SR,Ca,ina}} + \mathrm{Ca_i}} \cdot \frac{\mathrm{Ca_i}}{\mathrm{Ca_i} + K_{\mathrm{SR,Ca,act}}} \cdot \frac{\mathrm{IP_3}}{\mathrm{IP_3} + K_{\mathrm{SR,IP3}}}, \tag{S25}$$

$$J_{\mathrm{IP3R}} = J_{\mathrm{IP3R,max}}\,P_{\mathrm{IP3R}}^3\,(\mathrm{Ca_s} - \mathrm{Ca_i}), \tag{S26}$$

$$I_{\mathrm{SERCA}} = I_{\mathrm{SERCA,max}} H(\mathrm{Ca_i}; n_{\mathrm{SERCA}}, K_{\mathrm{SERCA}}). \tag{S27}$$

The remaining transporter/pump currents are

$$I_{\mathrm{PMCA}} = I_{\mathrm{PMCA,max}}\,H(\mathrm{Ca_i};\, n_{\mathrm{PMCA}}, K_{\mathrm{PMCA}}), \tag{S28}$$

$$I_{\mathrm{NaK}} = I_{\mathrm{NaK,max}}\,H(\mathrm{K_o};\, n_{\mathrm{NaK,Ko}}, K_{\mathrm{NaK,Ko}})\,H(\mathrm{Na_i};\, n_{\mathrm{NaK,Nai}}, K_{\mathrm{NaK,Nai}})\,\frac{V_{\mathrm{m}} + 150}{V_{\mathrm{m}} + 200}, \tag{S29}$$

The $Na^+/Ca^{2+}$ exchanger current is

$$I_{\mathrm{NCX}} = g_{\mathrm{NCX}}\,\frac{\mathrm{Na_i^3 Ca_o}\,\Phi_F - \mathrm{Na_o^3 Ca_i}\,\Phi_R}{1 + d_{\mathrm{NCX}}\left(\mathrm{Na_o^3 Ca_i} + \mathrm{Na_i^3 Ca_o}\right)}, \tag{S30}$$

with

$$\Phi_F = \exp\!\left(\frac{\gamma_{\mathrm{NCX}} F V_{\mathrm{m}}}{RT}\right), \qquad \Phi_R = \exp\!\left(\frac{(1-\gamma_{\mathrm{NCX}}) F V_{\mathrm{m}}}{RT}\right). \tag{S31}$$

Finally, the $Na^+$-$K^+$-$2Cl^-$ cotransporter current is

$$I_{\mathrm{NKCC1}} = L_{\mathrm{NKCC1}}\,\ln\!\left(\frac{\mathrm{K_o Na_o Cl_o^2}}{\mathrm{K_i Na_i Cl_i^2}}\right)\frac{K_{\mathrm{NKCC1,ina}}}{K_{\mathrm{NKCC1,ina}} + \mathrm{Cl_i}}. \tag{S32}$$

### S4 Objective-Function Definition

The residual is built from four distinct building blocks: (i) a one-sided band-violation penalty, (ii) a steady-state convergence residual, (iii) relative-error terms enforcing current-ordering constraints, and (iv) seed-robustness penalty.

**Band-violation penalty** For a quantity $x$ constrained to an admissible band $[x_1, x_2]$ (a physiological range, or an experimental measurement $\times(1 \pm r_{\mathrm{err}})$), we define the squared deviation from the nearest violated bound as:

$$e_{\mathrm{b}}(x; x_1, x_2) = \begin{cases} (x_1 - x)^2, & x < x_1 \\ (x - x_2)^2, & x > x_2 \\ 0, & x_1 \leq x \leq x_2 \end{cases} \tag{S33}$$

and the associated relative error is defined as $\epsilon_{\mathrm{b}}(x; x_1, x_2) = e_{\mathrm{b}}(x; x_1, x_2)/x_m^2$ where $x_m = (x_1 + x_2)/2$, except when the midpoint is (effectively) zero, in which case $x_m = x_2 - x_1$. Both forms vanish whenever $x$ lies inside $[x_1, x_2]$ and grow with the distance to the nearest violated bound.

**Steady-state convergence residual** For a candidate equilibrium $\mathbf{x}^*(\sigma)$ obtained by integrating the ODE system to steady state at pressure $\sigma$:

$$\epsilon_{\mathrm{r}}(\mathbf{x}^*, \sigma) = \frac{1}{N} \sqrt{\sum_{k=1}^{N} \dot{x}_k\big(\mathbf{x}^*, \sigma\big)^2}\,, \tag{S34}$$

with $\dot{\mathbf{x}} \to 10^6 \dot{\mathbf{x}}$ applied beforehand whenever $\max_k |\dot{x}_k(\mathbf{x}^*, \sigma)| > 10^{-6}$. This term is a numerical convergence check (not a fit to data), penalizing states that are not genuine equilibria of the governing ODEs.

**Current-ordering penalty** Considering two generic currents (at rest) $I_i$ and $I_j$, we define the current-ordering (squared relative) error as

$$\varepsilon_{\mathrm{curr}} = \left( \frac{|I_i| - |I_j|}{|I_j|} \right)^2 \mathbb{1}\{|I_i| > |I_j|\}. \tag{S35}$$

**Seed-robustness penalty** For a fixed probe set $\mathcal{S}$ perturbing the initial condition, $\mathbf{x}_0 \to \mathbf{x}_0 \odot (1 + \boldsymbol{\delta}_s)$, $\boldsymbol{\delta}_s \sim \mathcal{U}(-p, p)^N$, $p = 0.05$:

$$\varepsilon_{\mathrm{seed}}(\sigma) = k_{\mathrm{seed}} \max_{s \in \mathcal{S}} e\big(V_m^{(s)};\, 0.99\, V_m^*,\, 1.01\, V_m^*\big)\,, \qquad k_{\mathrm{seed}} = 10^3 \tag{S36}$$

where $V_{\mathrm{m}}^{(s)}$ is the equilibrium reached from the $s$-th perturbed seed and $V_{\mathrm{m}}^*$ the unperturbed reference. Unlike the constraints above, this term uses the unnormalized deviation $e_{\mathrm{b}}(\cdot)$ directly (acting as a tolerance/bistability trigger), and a large penalty activates as soon as any probe seed converges to a different branch of the (potentially multistable) system.
